# Mismatch repair deficiency is not sufficient to increase tumor immunogenicity

**DOI:** 10.1101/2021.08.24.457572

**Authors:** Peter M K Westcott, Francesc Muyas, Olivia Smith, Haley Hauck, Nathan J Sacks, Zackery A Ely, Alex M Jaeger, William M Rideout, Arjun Bhutkar, Daniel Zhang, Mary C Beytagh, David A Canner, Roderick T Bronson, Santiago Naranjo, Abbey Jin, JJ Patten, Amanda M Cruz, Isidro Cortes-Ciriano, Tyler Jacks

## Abstract

DNA mismatch repair deficiency (MMRd) in human cancer is associated with high tumor mutational burden (TMB), frameshift mutation-derived neoantigens, increased T cell infiltration, and remarkable responsiveness to immune checkpoint blockade (ICB) therapy. Nevertheless, about half of MMRd tumors do not respond to ICB for unclear reasons. While tumor cell line transplant models of MMRd have reinforced the importance of TMB in immune response, critical questions remain regarding the role of immunosurveillance in the evolution of MMRd tumors induced *in vivo*. Here, we developed autochthonous mouse models of lung and colon cancer with highly efficient ablation of MMR genes via *in vivo* CRISPR/Cas9 targeting. Surprisingly, MMRd in these models did not result in increased immunogenicity or response to ICB. Mechanistically, we showed this lack of immunogenicity to be driven by profound intratumoral heterogeneity (ITH). Studies in animals depleted of T cells further demonstrated that immunosurveillance in MMRd tumors has no impact on TMB but shapes the clonal architecture of neoantigens by exacerbating ITH. These results provide important context for understanding immune evasion in cancers with high TMB and have major implications for therapies aimed at increasing TMB.

## Introduction

ICB therapies have revolutionized the treatment landscape for a range of tumor types, particularly those with high TMB^1–4^. Somatic protein-altering (non-synonymous) mutations in the cancer genome can generate novel antigens (neoantigens) that are presented by major histocompatibility complexes (MHC) and elicit tumor-specific T cell responses^5,6^. It is therefore widely accepted that increased TMB, and by extension neoantigen burden, renders tumors susceptible to T cell attack following re-invigoration by immunotherapy. A number of studies have demonstrated such a correlation and proposed that TMB may be a valuable clinical biomarker for identifying patients that will respond favorably to ICB within and across tumor types^7–9^. In particular, MMRd is associated with some of the highest mutational burdens observed in cancer^10–13^, termed hypermutation. A clinical trial of pembrolizumab (anti-PD-1) across MMRd tumors spanning 12 cancer types demonstrated remarkable efficacy^3^, leading to pan-cancer FDA approval for this indication. More recently, FDA approval was granted to pembrolizumab for all tumors based on high TMB alone^8^.

Nevertheless, about half of patients with MMRd tumors did not respond to pembrolizumab, and TMB was not predictive of response in these patients^3^, underscoring the critical need to understand what factors beyond MMRd and TMB mediate response to immunotherapy. Interestingly, a number of previously reported correlations of TMB with ICB response did not withstand reanalysis incorporating rigorous statistical corrections for important covariates like tumor subtype^14^. A recent study argued that FDA approval of pembrolizumab based on TMB may be too broad, overlooking important covariates that underlie both favorable response to ICB and increased TMB, such as the initiating carcinogen^15^. Finally, while a T cell inflamed tumor microenvironment is one of the most robust predictors of ICB response, TMB is poorly correlated with degree of T cell infiltration in pan-cancer analyses^16,17^.

One likely reason for the weak correlation of TMB with ICB response is intratumoral heterogeneity (ITH) of mutations. Neoantigens of low tumor cellularity, or clonality, could presumably be deleted with minimal impact on overall tumor fitness^18^, or fail to elicit productive T cell responses altogether^19^. Indeed, it has been observed in a number of human cancers that ITH is associated with decreased T cell infiltration^20,21^ and poor survival^22,23^, while clonal neoantigens are predictive of better response to ICB^23,24^. This concept has been exemplified in an experimental setting of UVB-induced hypermutation of melanoma cell lines followed by single cell cloning and retransplant into immune-competent mice^25^. This study found that low heterogeneity and diversity of mutations across tumor clones resulted in greater immunogenicity and response to ICB^25^. It is thus reasonable to hypothesize that similar mechanisms are operative in MMRd tumors, especially given their constitutive mutational instability. Surprisingly, the role of ITH in the evolution and immune evasion of MMRd tumors remains poorly understood.

Deletion of *Mlh1*, an essential component of the MMR complex, in cancer cell lines *in vitro* has been shown to cause mutational instability and increased responsiveness to immunotherapy^26,27^, with prolonged growth in culture corresponding to greater immunogenicity. However, a major caveat of these studies is the accumulation of mutations *in vitro* in the absence of immunosurveillance. Importantly, it is well-appreciated that tumor immunogenicity, or lack thereof, is exquisitely shaped by crosstalk with the immune system, a process termed immunoediting^5,28,29^. It is however unclear how the heterogeneous and continuously evolving landscape of neoantigens in MMRd tumors is shaped in an immune-competent host, and whether induction of MMRd *in vivo* is sufficient to render tumors immunogenic or responsive to ICB.

To specifically address the role of mutational instability in ITH and tumor immunogenicity in a physiologically relevant setting, we developed autochthonous mouse models of MMRd lung and colon cancer by disrupting the MMR pathway genes *Msh2*, *Mlh1*, *Msh3*, and *Msh6*. We found that MMRd in our models was not sufficient to render lung or colon tumors immunogenic or responsive to ICB. As we describe below, sustained mutational instability drove profound ITH, with immunosurveillance sculpting the clonal architecture, but not overall burden, of neoantigens.

## Results

### *In vivo* cancer models with CRISPR/Cas9-targeted disruption of MMR show high mutational burden

We first adapted the autochthonous mouse model of lung cancer developed in our laboratory^30^ by breeding a *Cas9*-expressing allele with Cre-inducible oncogenic *Kras* and *Trp53* loss-of-function alleles (*R26^LSL-Cas9^; Kras^LSL-G12D^*; *Trp53^flox/flox^*). Intratracheal instillation of lentivirus expressing Cre and an *Msh2*-targeting sgRNA (sgMsh2) into these animals resulted in lung adenocarcinomas with highly efficient protein knockout of MSH2 (∼80% of week 16 tumor area) (Fig. 1a-c, Supplemental Fig. 1a-b). Multiple cell lines generated from these tumors confirmed loss of MSH2 protein and frameshifting insertions/deletions (indels) at the targeted locus, while some lines harbored in-frame edits and retained protein expression (Supplemental Fig. 1c-d). We also adapted an endoscope-guided submucosal injection technique^31^ to deliver lentivirus carrying sgRNAs targeting the essential colon tumor suppressor, *Apc*, in tandem with *Msh2*, *Mlh1*, *Msh3*, or *Msh6*, into the distal colon of mice with constitutive *Cas9* expression (Fig. 1b). This resulted in highly efficient induction of focal colon adenomas with MMR gene knockout (Fig. 1d, Supplemental Fig. 1e-i).

**Figure 1.**
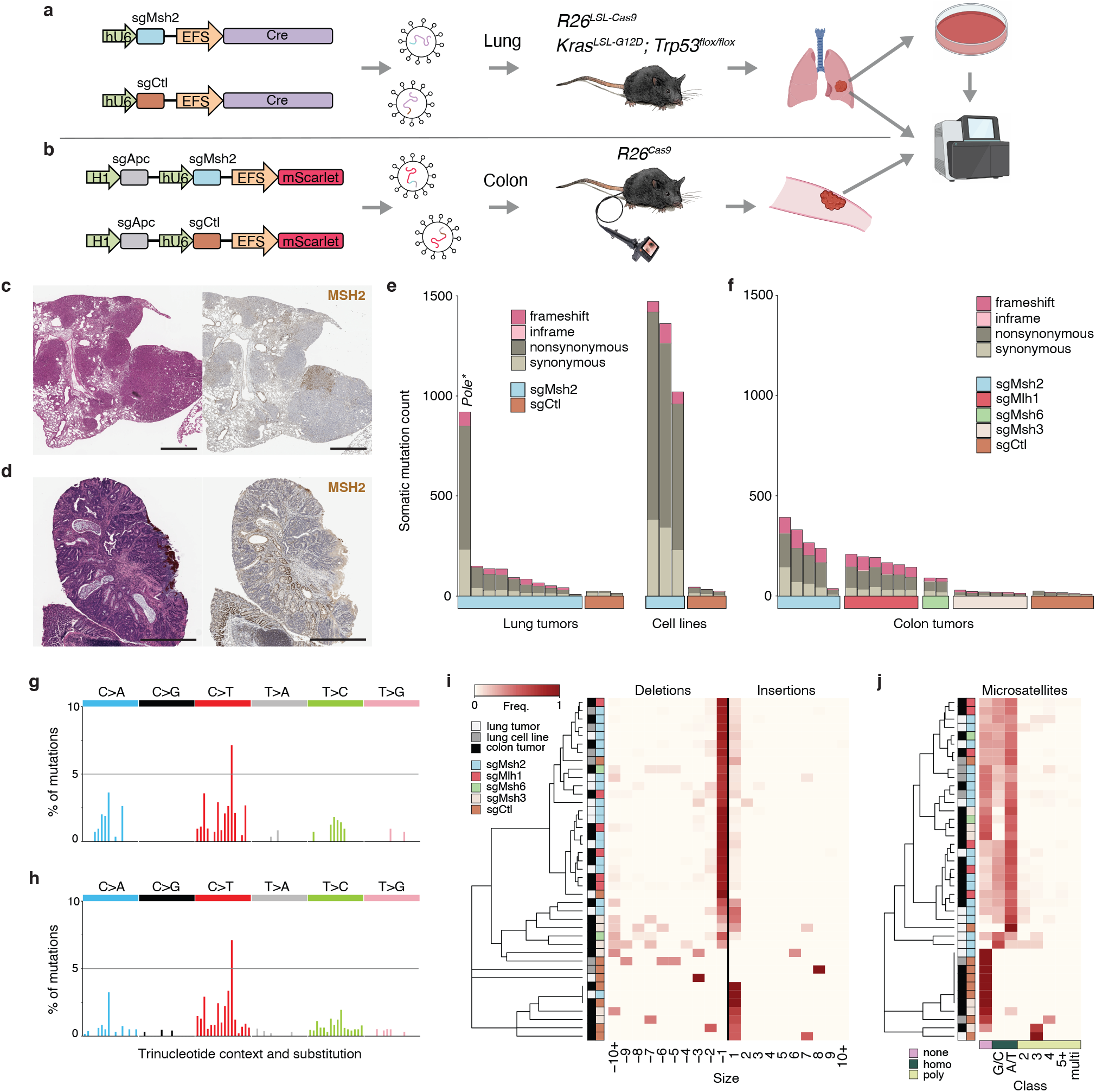
Development of *in vivo* CRISPR/Cas9-targeted models of MMRd lung and colon cancer. **(a-b)** Schematic of lentiviral constructs and mouse strains used to induce MMRd lung **(a)** and colon **(b)** tumors for WES and *in vitro* analyses. **(c-d)** H&E and MSH2 IHC of sgMsh2-targeted lung **(c)** and colon **(d)** tumors 16-weeks post-initiation, representative of 10 independent animals each. Scale bars = 1 mm. **(e-f)** Total somatic SNVs (grey shades) and indels (pink shades) identified in the protein coding exome of autochthonous lung tumors and cell lines **(e)** and autochthonous colon tumors **(f)**. *Pole\** = *Pole* S415R mutation. **(g-h)** Median percentage of the 96 possible SNVs classified by substitution and flanking 5’ and 3’ bases observed across 9 sgMsh2-targeted lung tumors (excluding one tumor with *Pole* S415R mutation) **(g)** and 5 sgMsh2-targeted colon tumors **(h)**. **(i-j)** Frequency of indels from -10 to 10 nucleotides **(i)** and the frequency of indels occurring in different DNA microsatellite contexts **(j)** across all sequenced autochthonous tumors and parental cell lines. In **(j)**, none = no microsatellite, homo = homopolymer runs of four or more G/C or A/T bases, 2-5+ = microsatellites with motifs of 2-5+ bases, and multi = microsatellites with multiple repetitive motifs.

To confirm mutation of MMR genes and investigate the degree of TMB, we performed whole-exome sequencing (WES) on 37 micro-dissected tumors at 16-20 weeks post initiation, including 10 sgMsh2- and 3 non-targeting control (sgCtl)-targeted lung tumors, and 5 sgMsh2-, 6 sgMlh1-, 2 sgMsh6-, 6 sgMsh3-, and 5 sgCtl-targeted colon tumors. Targeted amplicon sequencing of the sgRNA-targeted MMR gene loci in these tumors confirmed extensive frame-shifting indels, indicating successful gene knockout. sgMsh2-, sgMlh1-, and sgMsh6-targeted lung and colon tumors showed an increased burden of somatic single nucleotide variants (SNVs) and indels, while sgMsh3-targeted colon tumors showed an increased burden of indels but not SNVs (Fig. 1e-f, Supplemental Fig. 1j). Mutational spectra of MMR gene-targeted lung and colon tumors were highly consistent with those observed in human MMRd tumors^32^. Specifically, SNVs were mostly C>T transitions and indels mostly single nucleotide deletions enriched at A/T homopolymer microsatellite repeats (Fig. 1g-j, Supplemental Fig. 1k-l). Consistent with the specific role of *Msh3* in the repair of large insertion-deletion loops, sgMsh3-targeted colon tumors showed relatively more large deletions, which were not enriched at microsatellites (Fig. 1i-j). Interestingly, one sgMsh2-targeted lung tumor had an S415R mutation in the exonuclease domain of DNA polymerase epsilon (*Pole*) and a much higher TMB (Fig. 1e, Supplemental Fig. 1m), consistent with the “ultramutator” phenotype of polymerase exonuclease domain mutant tumors in mouse and human^33,34^. These results establish the utility of our *in vivo* tumor models to recapitulate fundamental mutational features of MMRd in human cancer.

We also performed WES on 6 cell lines derived from *in vivo* sgMsh2- and sgCtl-targeted lung tumors. Unexpectedly, sgMsh2-targeted cell lines showed a 10-fold greater median TMB than comparable lung tumors (Supplemental Fig. 1j), suggesting either low purity of micro-dissected tumors, continued mutagenesis in culture, or substantial ITH that is reduced by clonal selection on plastic. Low tumor purity is likely only a minor contributor, as all lung tumors showed >78% purity based on recombination at the *Trp53^flox^* locus (see Methods). Similarly, the higher TMB we observed is unlikely a consequence of mutations acquired *in vitro* given the low number of passages. Consistent with high ITH that is reduced by *in vitro* culture, WES of a single cell clone (SSC) revealed more than double the TMB of the parental sgMsh2-targeted cell line, despite a relatively much smaller increase in TMB following 20 passages over two months in culture (Supplemental Fig. 1n). Furthermore, sequencing reads supporting 905 mutations private to the SSC (i.e., not called by our WES analysis pipeline in the parental line) were found in the sequencing data of the parental line, albeit at extremely low variant allele frequencies below threshold of mutation calling (Supplemental Fig. 1o). While some of these mutations could be sequencing errors, the same analysis in an unrelated control line (13-1) found 5.5-fold fewer mutations (Supplemental Fig. 1p). A similar disparity between mutation calling in bulk versus single-cell DNA sequencing was previously observed in hypermutated gliomas^35^.

Altogether, these results establish that our models recapitulate the mutational processes underlying hypermutation in MMRd human cancer. The lower clonal TMB we observed is likely due to neutral evolution in the absence of major selective bottlenecks following initiation. Given that ITH is associated with more aggressive disease and decreased response to ICB in humans^22–24^, our models are uniquely suited to study the role of ITH in immune dysfunction of cancers with high (colon) and low (lung) prevalence of MMRd alike.

### MMRd does not increase T cell infiltration or response to immune checkpoint blockade

Next, we sought to determine the effects of MMRd on the kinetics of tumorigenesis and T cell infiltration in our models of lung and colon cancer. To provide orthogonal validation of our CRISPR/Cas9 MMRd lung model, we also used mice harboring a Cre-inducible knockout allele with loxP sites flanking exon 12 of *Msh2*^36^ (*Kras^LSL-G12D^*; *Trp53^flox/flox^*; *Msh2^flox/flox^*—KPM). KPM and MMR proficient control (*Kras^LSL-G12D^*; *Trp53^flox/flox^*—KP) tumors were induced by intratracheal instillation of adenovirus expressing Cre driven by the alveolar type II cell-specific surfactant protein C (SPC) promoter. First, we took whole lungs at 5- or 15-16-weeks post-initiation and performed histological and immunohistochemical analyses. MSH2 protein knockout in the sgMsh2-targeted and KPM models was 79% and 89% efficient in late-stage lung tumors, respectively (Supplemental Figs. 1a, 2a). Interestingly, the percent area of tumors with MSH2 knockout in the CRISPR/Cas9 model significantly increased from 5 to 16 weeks (Supplemental Fig. 1a), suggesting positive selection. However, there was no significant difference in overall tumor burden or tumor grade with *Msh2* targeting at either time point in either model (Fig. 2a-d, Supplemental Fig. 2b-c). To our surprise, there was also no significant difference in tumor infiltration by T cells (CD3^+^) in the sgMsh2-targeted CRISPR/Cas9 model (Supplemental Fig. 2d-e), or whole lung infiltration by cytotoxic (CD8^+^), helper (CD4^+^), or regulatory (CD4^+^/FOXP3^+^) T cells in the KPM model (Fig. 2e-g), at either time point. This contrasts with a recent study that reported increased T cell infiltration in KPM lung tumors at 4-weeks post-initiation^37^. However, fewer KPM animals were analyzed in this study (N = 4), and statistical significance was not assessed at the whole lung level by animal, as we have done. Next, we assessed the effects of *Msh2* targeting on colon tumorigenesis in our colonoscopy guided CRISPR/Cas9 model. Like the lung models, there was no significant difference in growth kinetics of sgCtl-versus sgMsh2-targeted colon tumors, as measured by longitudinal colonoscopy (Supplemental Fig. 2f).

**Figure 2.**
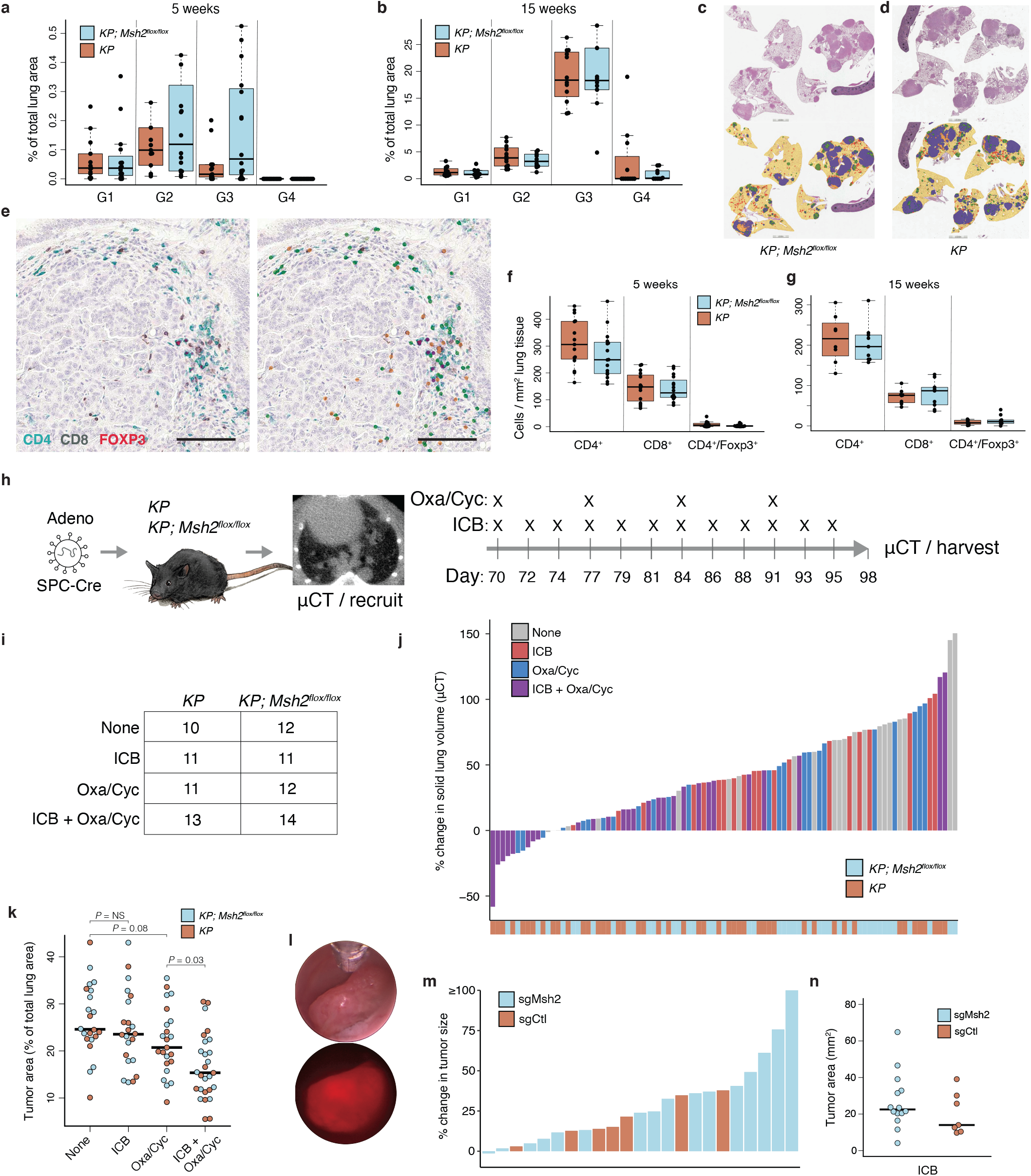
MMRd models of lung and colon cancer are not immunogenic. **(a-b)** Percent of total lung area occupied by tumors of grades 1-4 (G1-4) in *KP; Msh2^flox/flox^* and *KP* models at 5- **(a)** and 15-weeks **(b)** post-initiation with SPC-Cre adenovirus, representative of 16 and 15 animals at 5-weeks and 10 and 12 animals at 15-weeks. Normal lung and tumor grades were quantified using an automated convoluted neural network (CNN) developed in collaboration with Aiforia. **(c-d)** Whole tumor-bearing lung H&E and CNN annotation masks from *KP; Msh2^flox/flox^* **(c)** and *KP* **(d)** models 15-weeks post-initiation, representative of animals in (b). Yellow = normal lung; red = G1; green = G2; blue = G3; orange = G4. **(e)** IHC staining of lung tumor from 15-week *KP; Msh2^flox/flox^* mouse, representative of 10 animals (left). Green = CD4; black = CD8α; red = FOXP3. CD4^+^, CD8^+^, and regulatory T cell annotation masks generated by an automated CNN developed in collaboration with Aiforia (right). Scale bars = 100 μM. **(f-g)** Aiforia CNN quantification of CD4^+^, CD8^+^, and regulatory (CD4^+^/FOXP3^+^) T cell numbers in tumor-bearing lungs of *KP; Msh2^flox/flox^* and *KP* models at 5- **(f)** and 15-weeks **(g)** post-initiation, representative of 16 and 15 animals at 5-weeks and 9 and 8 animals at 15- weeks. **(h-i)** Schematic of preclinical trial design in *KP; Msh2^flox/flox^* and *KP* models **(h)**, and number of animals in treatment arms **(i)**. **(j)** Change in solid lung volume as measured by μCT pre-treatment (10 weeks) and post-treatment (14 weeks). **(k)** Lung tumor burden at necropsy (14 weeks) as measured by manual annotation of H&E-stained whole lung sections. **(l)** Brightfield and fluorescent colonoscopy images of sgMsh2-targeted colon tumor, representative of 16 animals. Reproducible placement of biopsy forceps in brightfield image allows normalization of relative tumor area. **(m)** Change in colon tumor size by colonoscopy pre-treatment (20 weeks) and post-treatment (24 weeks). N = 16 sgMsh2- and 7 sgCtl-targeted animals. **(n)** Colon tumor burden at necropsy (24 weeks) as measured by stereomicroscope image. N = 14 sgMsh2- and 7 sgCtl-targeted animals. Significance in (k) was assessed by Wilcoxon Rank Sum test.

To determine the sensitivity of our models to immunotherapy, we first performed preclinical trials with ICB (combined anti-CTLA-4/PD-1) in the KPM lung model. Additionally, we included treatment with the chemotherapeutic combination of oxaliplatin and low-dose cyclophosphamide (Oxa/Cyc) alone and in combination with ICB, as Oxa/Cyc has previously been shown to induce immunogenic cell death and synergize with ICB in a variant of the KP lung cancer model harboring a model neoantigen^38,39^. Mice were randomly enrolled into treatment arms following confirmation of discrete lung tumors by X-ray microcomputed tomography (μCT) at 10 weeks post-initiation and dosed for one month (Fig. 2h-i). At treatment conclusion, mice were imaged again by μCT and necropsied for histological quantification of lung tumor burden. To our surprise again, no significant differences were observed between KPM and KP mice across all treatment arms, both in longitudinal change (Fig. 2j) or final tumor burden at necropsy (Fig. 2k). Interestingly, the combination of Oxa/Cyc and ICB reduced tumor burden significantly more than Oxa/Cyc or ICB alone in both KPM and KP mice and to a similar extent (Fig. 2k). These results were not unique to the lung, as ICB treatment failed to induce any responses (as defined by Response Evaluation Criteria in Solid Tumors) in sgMsh2-targeted colon tumors (Fig. 2l-m, Supplemental Fig. 2g-h). Likewise, there was no significant difference in endpoint size of sgCtl-versus sgMsh2-targeted colon tumors following ICB treatment (Fig. 2n). Altogether, these data suggest that MMRd is not sufficient to increase sensitivity of tumors to immunotherapy, in stark contrast to previous reports^26,27^.

### Somatic loss of MMR drives profound intratumoral heterogeneity of mutations

To assess the clonal composition of mutations in our models, we first estimated the cancer cell fraction (CCF) of mutations using a previously published algorithm^40^. This revealed that most mutations were present in less than a quarter of cells (CCF = 0.25) in sgMsh2-targeted lung tumors and cell lines and of MMR gene-targeted colon tumors (Fig. 3a-b, Supplemental Fig. 3a). To investigate the extent of ITH more deeply, we isolated and performed WES on 8 SSCs derived from an MSH2 knockout cell line (09-2). Prior to subcloning 09-2, we restored MMR by re-expressing *Msh2* using a bicistronic lentivirus with the puromycin resistance gene (Fig. 3c). Importantly, SSCs derived from this line (M1-8) maintained MSH2 protein expression and showed minimal accumulation of additional mutations and stable clonal architecture after 20 passages in the presence of puromycin over two months (Fig. 3d, Supplemental Fig. 3b-c). Despite this mutational stability, significantly more somatic mutations were called in all SSCs than the MSH2 knockout parental lines (Fig. 3e), further supporting the notion that the degree of ITH is underestimated by bulk sequencing methods^35^. To assess the clonal representation of the SSCs in the parental line from which they were derived, we identified four unique SNVs in copy number neutral regions for each SSC and performed ultradeep targeted amplicon sequencing of the parental line (see Methods). This identified three of the SSCs above background, only one of which was greater than 1% clonal fraction of the parental line (Fig. 3f). To determine clonal relationships between SSCs, we compared mutation overlap and constructed a phylogenetic tree rooted on the parental line (Fig. 3g, Supplemental Fig. 3d). This further highlighted extensive dissimilarity of mutations across clones and a staggering level of ITH.

**Figure 3.**
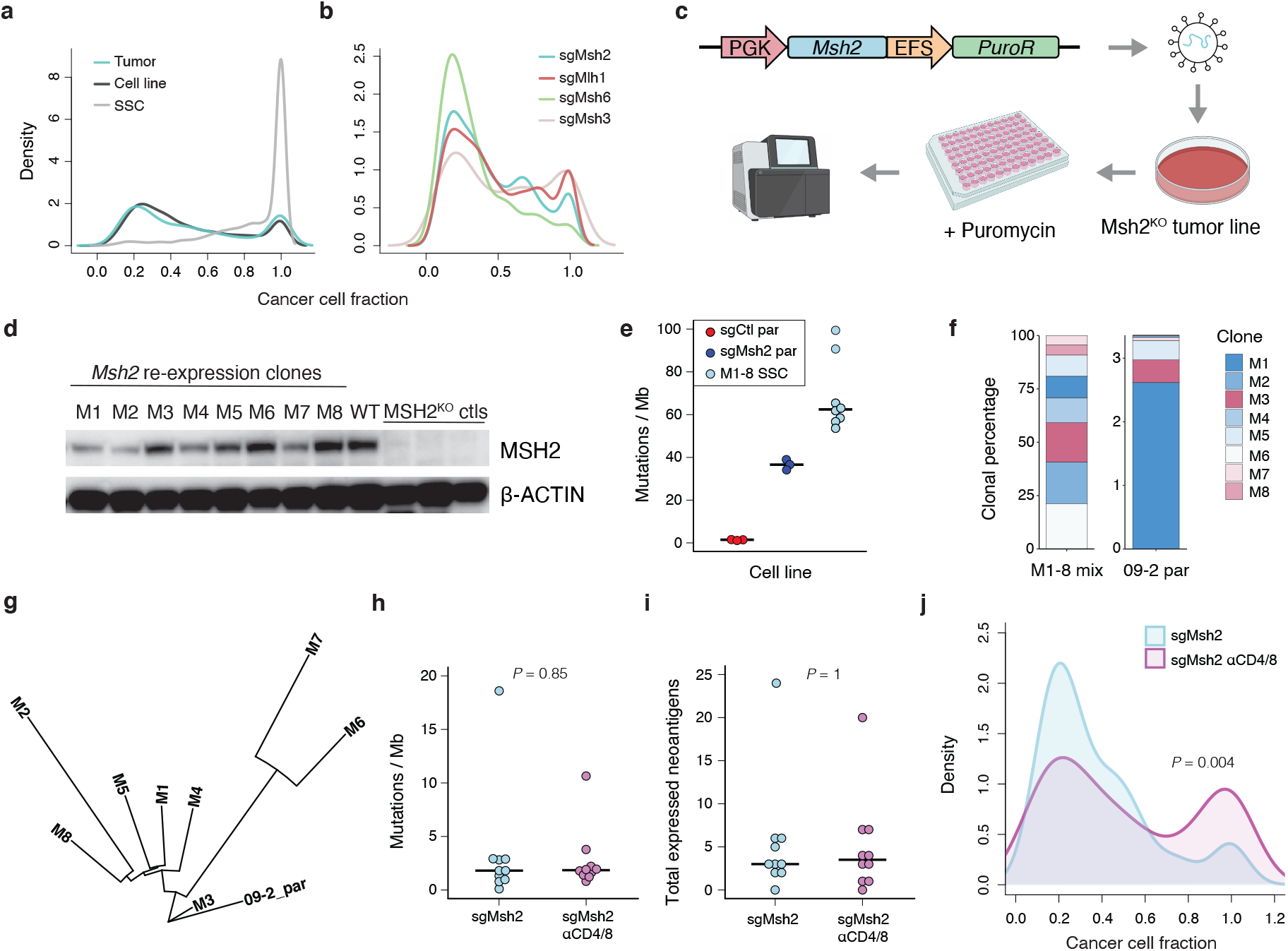
MMRd models are defined by extensive ITH. **(a-b)** Distribution of cancer cell fraction estimates of all SNVs in lung tumors, cell lines, and SSCs M1-8 **(a)** and colon tumors **(b)**. Smoothing was performed by Gaussian kernel density estimation. **(c)** Schematic of single cell cloning workflow with re-expression of *Msh2*. **(d)** Western blot of MSH2 expression in SSCs with *Msh2* re-expression. WT = sgCtl-targeted cell line; MSH2^KO^ ctls = parental sgMsh2-targeted cell lines. **(e)** Total mutations identified in *ex vivo* lung tumor-derived cell lines and SSCs, as mutations / Mb of DNA. Par = parental cell line. N = 3 sgCtl, 3 sgMsh2, and 8 M1-8 SSC lines. **(f)** Estimation of clonal frequencies of M1-8 SSCs in equimolar mixture of SSC DNA (left) and 09-2 parental line (right), as determined by targeted deep amplicon sequencing of 4 private SNVs per SSC. **(g)** Phylogenetic tree of clonal relationships between M1-8 SSCs, rooted on the parental line 09-2 and constructed using shared mutations with the parsimonious ratchet method. **(h-i)** Total mutations / Mb **(h)** and predicted neoantigens with allele-specific expression > 0 **(i)** in 16–20-week sgMsh2-targeted autochthonous lung tumors from animals with and without continuous antibody-mediated T cell depletion (αCD4/8). N = 10 tumors per group. **(j)** Distribution of cancer cell fraction estimates of all SNVs in lung tumors from (h-i). Smoothing was performed by Gaussian kernel density estimation. Significance was assessed by two-sided Kolmogorov-Smirnov test. Significance in (h-i) was assessed by Wilcoxon Rank Sum test.

Given that ITH in our model arose *in vivo* in an immune-competent host, we next sought to understand the role of immunosurveillance in shaping this process. We performed WES on 10 micro-dissected sgMsh2-targeted lung tumors (week 16-20) from animals depleted of CD8^+^ and CD4^+^ T cells by continuous anti-CD8 and anti-CD4 antibody dosing. Interestingly, these tumors did not harbor a higher TMB than tumors from immune-competent animals (Fig. 3h). To focus on potentially immunogenic mutations only, we adapted a neoantigen prediction pipeline we previously developed (Westcott, et al, *Nature Cancer* 2021, in press)^41^ integrating multiple epitope affinity prediction algorithms^42–46^ for mouse H-2K^b^ and H-2D^b^ MHC-I alleles. We also performed RNA sequencing on matched tumor samples to determine allele-specific expression of neoantigens. Surprisingly, there was no evidence of increased burden of expressed predicted neoantigens (≤ 500 nM IC_50_ affinity) between tumors from immune-competent and depleted mice (Fig. 3i). However, there was a significant difference in the CCF distribution of these neoantigens, with an increased percentage of clonal neoantigens in tumors from T cell-depleted mice (Fig. 3j). This trend held even after removal of the one tumor from an immune-competent animal harboring a *Pole* S415R mutation, an outlier with the highest TMB and extensive ITH (Supplemental Fig. 3e). These results suggest that neoantigens with high, but not low, clonal fraction are negatively selected by the adaptive immune system. By extension, immunosurveillance in mutationally unstable tumors promotes ITH by increasing the overall fraction of subclonal neoantigens.

### Intratumoral heterogeneity promotes immune evasion and protects immunogenic subclones from deletion

To further evaluate the immunogenicity and ICB responsiveness of MMRd tumors in our model, we performed a series of survival experiments following orthotopic transplantation of the *ex vivo* tumor-derived cell lines and SSCs described above. Similar to previous reports of immunoediting shaping tumors to be non-immunogenic upon re-transplantation^5,28^, none of the parental MSH2 knockout lines resulted in accelerated disease with continuous T cell depletion, similar to the low TMB control line (Fig. 4a). Unlike MMRd in the autochthonous model, however, mice orthotopically transplanted with the parental line 09-2 showed 22% durable responses with ICB treatment (Fig. 4b). Transplant of an equal mixture of SSCs M1-8 resulted in modestly accelerated disease with T cell depletion and slightly better response to ICB, with 32% durable responses, suggesting increased immunogenicity compared to the parental line (Fig. 4c). Transplant of the SSCs individually revealed that some were strongly immunogenic (M3, M7, M8), forming tumors more aggressively with T cell depletion that resulted in significantly shortened survival (Fig. 4d-k). The other five SSCs did not form tumors more aggressively with T cell depletion, demonstrating that ITH is defined by both heterogeneity of mutations and immunogenic potential of subclones. These differences were not due to tumor intrinsic growth rates or defects in antigen presentation, as *in vitro* growth kinetics of SSCs were similar and did not correlate with *in vivo* aggressiveness, and all lines retained interferon gamma (IFNγ) sensitivity and MHC-I surface presentation (Supplemental Fig. 4a-e). Notably, two of the most aggressive and non-immunogenic clones (M1, M5) were highly responsive to ICB (Fig. 4l-m), suggesting that despite minimal baseline immunogenicity, clonal tumors are more sensitive to ICB treatment. Altogether, these results are consistent with decreasing ITH (autochthonous tumors > cell lines > SSC mixture > SSC) correlating with increased immunogenicity and ICB response, as has been observed in a flank transplant UVB mouse model^25^ and human cancer^24^.

**Figure 4.**
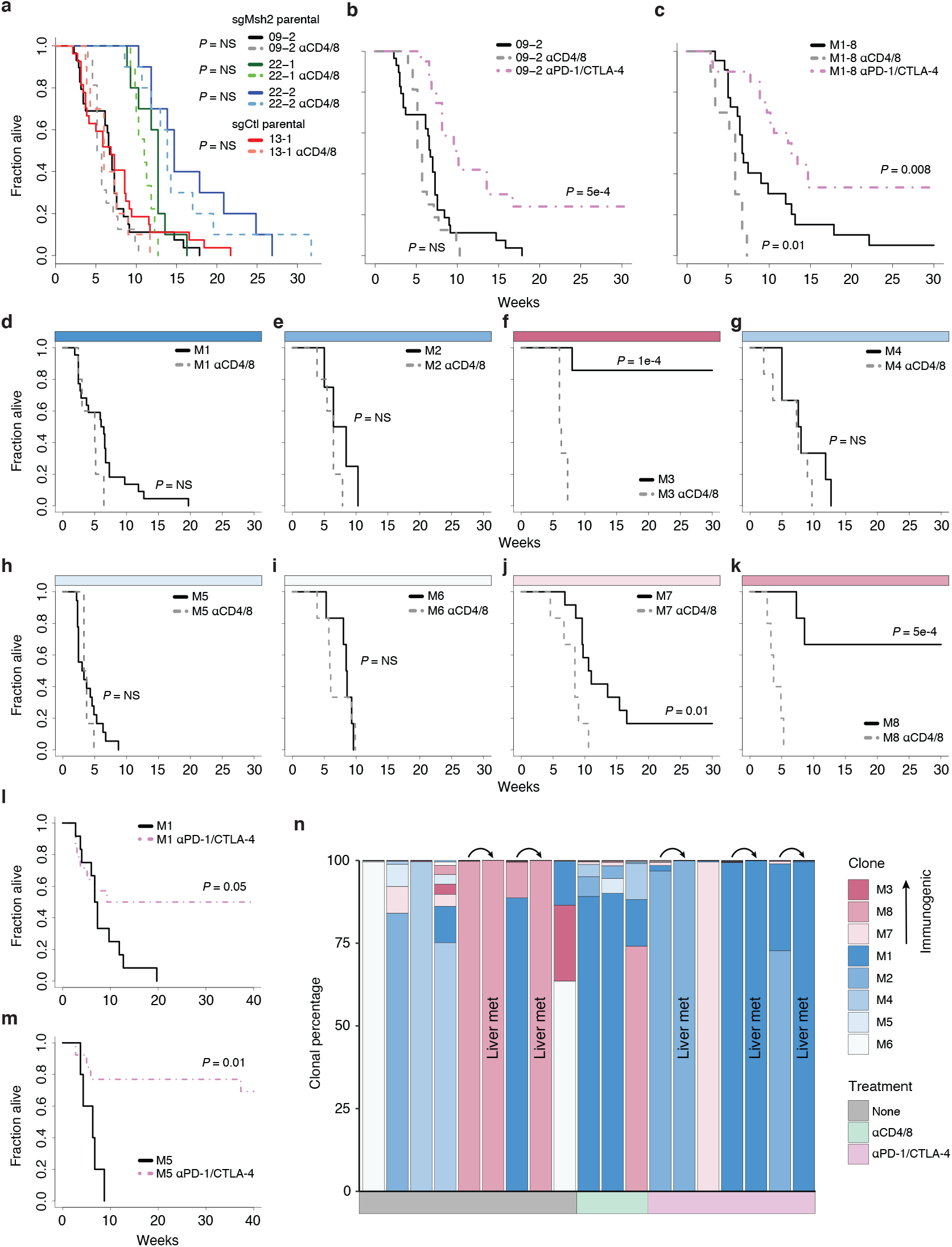
ITH promotes immune evasion of MMRd tumors. **(a-n)** Survival of syngeneic mice orthotopically transplanted via intratracheal instillation with indicated lung tumor cell lines and SSCs. **(a)** Transplant with parental sgMsh2- and sgCtl-targeted parental lines, with and without continuous T cell depletion (αCD4/8: lighter shades and dotted lines). **(b-c)** Transplant with parental 09-2 line **(b)** and equal mixture of M1-8 SSCs **(c)** with and without continuous αCD4/8, and ICB treatment. **(d-k)** Clonal transplants of M1-8 SSCs with and without continuous αCD4/8. **(l-m)** Transplant of M1 **(l)** and M5 **(m)** with and without ICB treatment. Mice that died before initiation of ICB treatment at 2 weeks were excluded from analysis in both arms. **(n)** Estimation of clonal percentages of M1-8 SSCs in lung tumors and liver metastases from animals transplanted with an equal mixture of all clones and receiving no treatment, continuous αCD4/8, or ICB. Tumors were collected from moribund mice over the course of the survival studies, between 10- and 20-weeks post-transplant for no treatment and ICB arms, and 5- and 7-weeks for αCD4/8 arm. Clonal percentages were determined by targeted deep amplicon sequencing of 4 private SNVs per SSC. Arrows above bars connect matched primary tumors and metastases. ICB treatment in (b-c) and (l-n) was started 2-weeks post-transplant and continued for 4 weeks. Significance in (a-m) was assessed by Cox proportional hazards regression, with Holm’s correction for multiple comparisons in (a) and (d-k).

The fact that the 09-2 cell line arose in an immune-competent host yet contained highly immunogenic subclones begs the question how did these clones originally evade immune deletion? We reasoned that ITH not only facilitates immune evasion by selective evolution of non-immunogenic clones but shields immunogenic subclones from deletion. To assess this hypothesis, we collected lung tumors and liver metastases from mice orthotopically transplanted with an equal mixture of M1-8 SSCs and reconstructed their clonal makeup by ultradeep targeted amplicon sequencing of SNVs private to each SSC (see Methods) (Fig. 4n). Despite the immunogenicity of M3, M7, and M8, these SSCs were detected in five of the seven mixed tumors analyzed from immune-competent animals at clonal fractions > 1%. Two of these animals also developed liver metastases, both of which were clonal outgrowths of M8. In contrast, none of the immune-competent animals transplanted with M3, M7, or M8 alone formed metastases. Therefore, in the face of competition with five non-immunogenic subclones, the immunogenic subclones persisted and, in the case of M8, grew out aggressively. This pattern was not substantially different than that of analogous transplants in T cell-depleted mice (Fig. 4n). However, ICB treatment resulted in selection against immunogenic subclones, with only one of four lung tumors harboring an immunogenic clone (M7) at > 1% fraction, and three liver metastases all derived from a single non-immunogenic subclone (Fig. 4n). Altogether, this analysis suggests that not all subclones within a tumor of high ITH are intrinsically immune evasive, but that subclones possess a range of potential immunogenicity that is exposed by ICB treatment. Importantly, sensitivity to ICB appears to be greatest at clonal purity and reduced with subclonality, consistent with previous studies in mouse and human^23–25^.

## Discussion

It is now well-established that somatic mutations can generate neoantigens capable of eliciting clinically meaningful anti-tumor immune responses^5,6^. Yet TMB, and thus neoantigen burden, is only modestly associated with response to immunotherapy^7–9,14,15^. About half of MMRd tumors do not respond to ICB, and TMB fails to stratify MMRd responders versus non-responders^3^. The mechanisms underlying these different responses must be elucidated before the benefits of immunotherapy can be expanded to more patients.

To address these questions, we developed autochthonous mouse models of lung and colon cancer with *in vivo* CRISPR/Cas9-targeted ablation of genes in the MMR pathway. These models recapitulate critical features of human lung and colon cancer, including genetics, histopathology, and *in situ* initiation in the relevant tissue microenvironment. Importantly, they enable study of mutations continuously acquired *in vivo* from tumor initiation through advanced disease. This is an important distinction, as multiple studies that experimentally demonstrated a role for MMRd and TMB in ICB response carried out mutagenesis *in vitro*^25–27^. Other studies employed tissue-specific knockout of *Msh2* or activation of dominant proofreading mutant *Pole* (*Pole^P286R^*) during embryogenesis to study the efficacy of prophylactic vaccination or ICB therapy on spontaneously arising hypermutant tumors in the mouse^47,48^. These models, which recapitulate familial cancers like Lynch syndrome in their accumulation of mutations in normal parenchyma preceding transformation, also showed response to immunotherapy.

In contrast, our models of MMRd lung and colon cancer did not display increased baseline immunogenicity or response to ICB, even when ICB was combined with an immunogenic chemotherapy regimen^39^ in the lung model. This definitively establishes that MMRd and mutational instability alone are not sufficient to increase tumor immunogenicity. The discrepancy of these results with past studies is likely due to timing of MMR inactivation and the resulting patterns of clonal expansion. In the present study, we induced MMRd concomitantly with tumor initiation, resulting in mutation accumulation during exponential cellular expansion. This is reminiscent of so called ‘born to be bad’ colon tumors that undergo a ‘Big Bang’ growth trajectory defined by the absence of selective sweeps^49^. Unlike models of familial cancer^47,48^, our models capture *de novo* loss of MMR in advanced tumors. Importantly, while Lynch syndrome accounts for less than 5% of CRC, about 10% of CRC harbors MMRd resulting from *MLH1* promoter hypermethylation or other somatic events^13^. Interestingly, it was recently shown that loss of MMR pathway activity in advanced gliomas occurring *de novo* or induced by alkylating chemotherapy treatment (e.g., temozolomide) led to extensive ITH and poor ICB response^35^.

Consistent with neutral evolution during the early stages of tumor progression and thus the absence of major clonal bottlenecks, tumors in our models showed profound ITH. Most mutations identified in autochthonous lung and colon tumors were present in less than a quarter of tumor cells. However, the full extent of ITH only became apparent upon WES of cell lines from *ex vivo Msh2* knockout lung tumors, which revealed an over 10-fold and nearly 20-fold increase in TMB in parental cell lines and SSCs, respectively. *Msh2* re-expression rendered cell lines mutationally static, and we provided multiple lines of evidence suggesting that these mutations were induced during *in vivo* tumorigenesis. Phylogenetic analysis of SSCs showed a low fraction of shared “truncal” mutations and substantial evolutionary divergence, suggesting that ITH began early in tumor evolution. Using deep amplicon sequencing of sets of private SNVs in each SSC, we showed that the most abundant clone comprised about 2.5% of the parental line, while all others comprised less than 1%. This extensive ITH and the inability to detect it by standard clinical sequencing depth has critical implications for the use of TMB as a biomarker. Given that differences in sequencing depth and analysis pipelines will greatly influence estimates of TMB, it is important for clinical pipelines to be standardized. Strategies to robustly assay ITH—a critical determinant of ICB response—such as multi-region or single-cell DNA sequencing, should enhance the predictive utility of TMB.

Consistent with previous studies in human and mouse^23–25^, we found that reconstitution of ITH by orthotopic re-transplantation of a mixture of SSCs into the lung potentiated immune evasion. Three of eight SSCs were highly immunogenic when transplanted alone, but poorly immunogenic when transplanted as a mixture with other clones, showing representation in mixed tumors and even seeding clonal metastases. What differentiates our study from a previous study of ITH induced by UVB mutagenesis^25^ is that mutations arose spontaneously *in vivo*. The fact that some of the SSCs we isolated are highly immunogenic suggests that they arose late in the context of an immunosuppressive microenvironment or were protected from deletion by high ITH in the original tumor. The results of our ITH reconstitution experiments strongly support the latter. One explanation for how ITH protects immunogenic subclones from deletion is low cellularity—in the case of seven of the eight SSCs, less than 1% of the parental cell line, at least at the four defining SNVs sequenced. This could result in insufficient material to cross prime T cell responses and early dysfunction or ignorance, as we and others have previously investigated (Westcott, et al, *Nature Cancer*, 2021, in press)^19^. Alternatively, ITH may result in a multitude of potentially immunogenic neoantigens, but the immune response is directed at a small subset of dominant antigens (Burger, et al., *Cell*, 2021, in press).

Paradoxically, we found that immunosurveillance may exacerbate ITH. Specifically, tumors from T cell-depleted mice showed no difference in TMB or burden of predicted neoantigens, but a significantly greater percentage of predicted neoantigens that were clonal. This argues that in the face of mutational instability, the immune system shapes the clonal architecture of tumors without driving deletion of most neoantigens. This result provides nuance to our understanding of immunoediting and highlights the advantage of mutationally unstable models like ours to study tumor evolution. Landmark studies that experimentally demonstrated immunoediting were performed in carcinogen models^5,28^, wherein a high burden of mutations is clonally fixed at initiation. These tumors are otherwise relatively mutationally static, restricting study of immunoediting of mutations to the earliest stages of tumorigenesis. In contrast, our models capture the continuous evolution of mutational instability and immunosurveillance throughout all stages of tumorigenesis, recapitulating a critical aspect of not only MMRd but the gradual accumulation of mutations seen in many human cancers^32^.

In summary, we developed new models of MMRd lung and colon cancer and showed that MMRd in the absence of clonal selection drives profound ITH that undermines the immune response. This raises important questions related to therapeutic strategies aimed at deliberately increasing TMB, which are being explored to increase tumor immunogenicity^26,50^. Our results argue that such strategies will likely fail to elicit meaningful immune engagement. More concerning, collateral mutagenesis that arises in this setting may drive more aggressive cancer, therapy resistance, or secondary malignancies. Future studies with models that enable temporal control of cooperating tumorigenic mutations will be helpful in determining the impact of clonal bottlenecks on immunosurveillance and immunotherapy response.

---

**Supplemental Figure 1.**
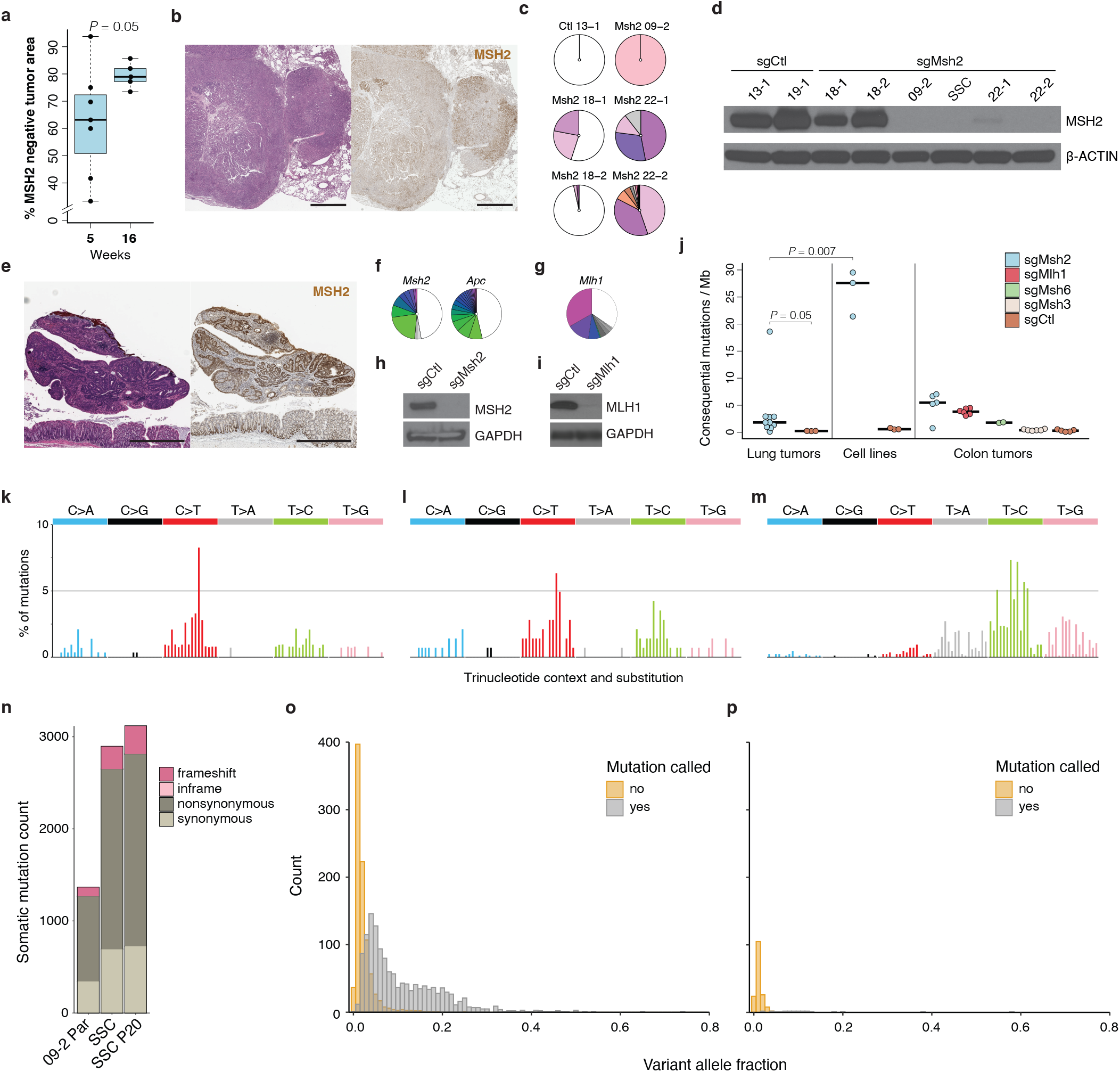
Validation of *in vivo* CRISPR/Cas9-targeted MMR gene knockout. **(a)** Percent of overall lung tumor area negative for MSH2 by IHC at 5- and 16-weeks post-initiation with sgMsh2 lentivirus. N = 7 animals at 5-weeks and 5 animals at 16-weeks. **(b)** H&E and MSH2 IHC of sgCtl (tdTomato)-targeted lung tumors at 16 weeks post-initiation. **(c)** Amplicon deep sequencing of sgMsh2-targeted locus in cell lines derived from an sgCtl- and five sgMsh2-targeted lung tumors harvested at 16-20 weeks post-initiation. White = unedited (wild-type); Colored = frameshifting mutations; Grey = inframe mutations. **(d)** Western blot of MSH2 protein expression in sgCtl- and sgMsh2-targeted lung tumor cell lines. SSC = single cell clone derived from 09-2 cell line. **(e)** H&E and MSH2 IHC of sgCtl (tdTomato)-targeted colon tumor at 16 weeks post-initiation, representative of 10 animals each. **(f-g)** Amplicon deep sequencing of sgMsh2/sgApc-targeted **(f)** and sgMlh1-targeted **(g)** loci in sgMsh2- and sgMlh1-targeted autochthonous colon tumors, representative of 5 sgMsh2- and 6 sgMlh1-targeted tumors. Colors same as (c). **(h-i)** Western blots of MSH2 **(h)** and MLH1 **(i)** protein expression in organoids derived from sgCtl-, sgMsh2-, and sgMlh1-targeted colon tumors, representative of one organoid line each. **(j)** Total consequential mutations (nonsynonymous SNVs and indels) for autochthonous lung tumors and cell lines and autochthonous colon tumors, shown as mutations / Mb of DNA. **(k-m)** Percentage of the 96 possible SNVs classified by substitution and flanking 5’ and 3’ base observed in a single sgMsh2-targeted lung tumor with *Pole* S415R mutation **(k)** and median percentage across 6 sgMlh1- **(l)** and 2 sgMsh6-targeted **(m)** colon tumors. **(n)** Total somatic SNVs (grey shades) and indels (pink shades) identified in 09-2 lung tumor cell line, early passage SSC derived from 09-2, and the same SSC after 20 passages. **(o-p)** Total number of mutations identified in early passage SSC that are also supported in sequencing reads of 09-2 parental line **(o)** and an unrelated MMR proficient control line, 13-1 **(p)**, with variant allele fraction on x-axis. Grey = mutations called by analysis pipeline; gold = mutations not called. Significance in (a) and (j) was assessed by Wilcoxon Rank Sum test.

**Supplemental Figure 2.**
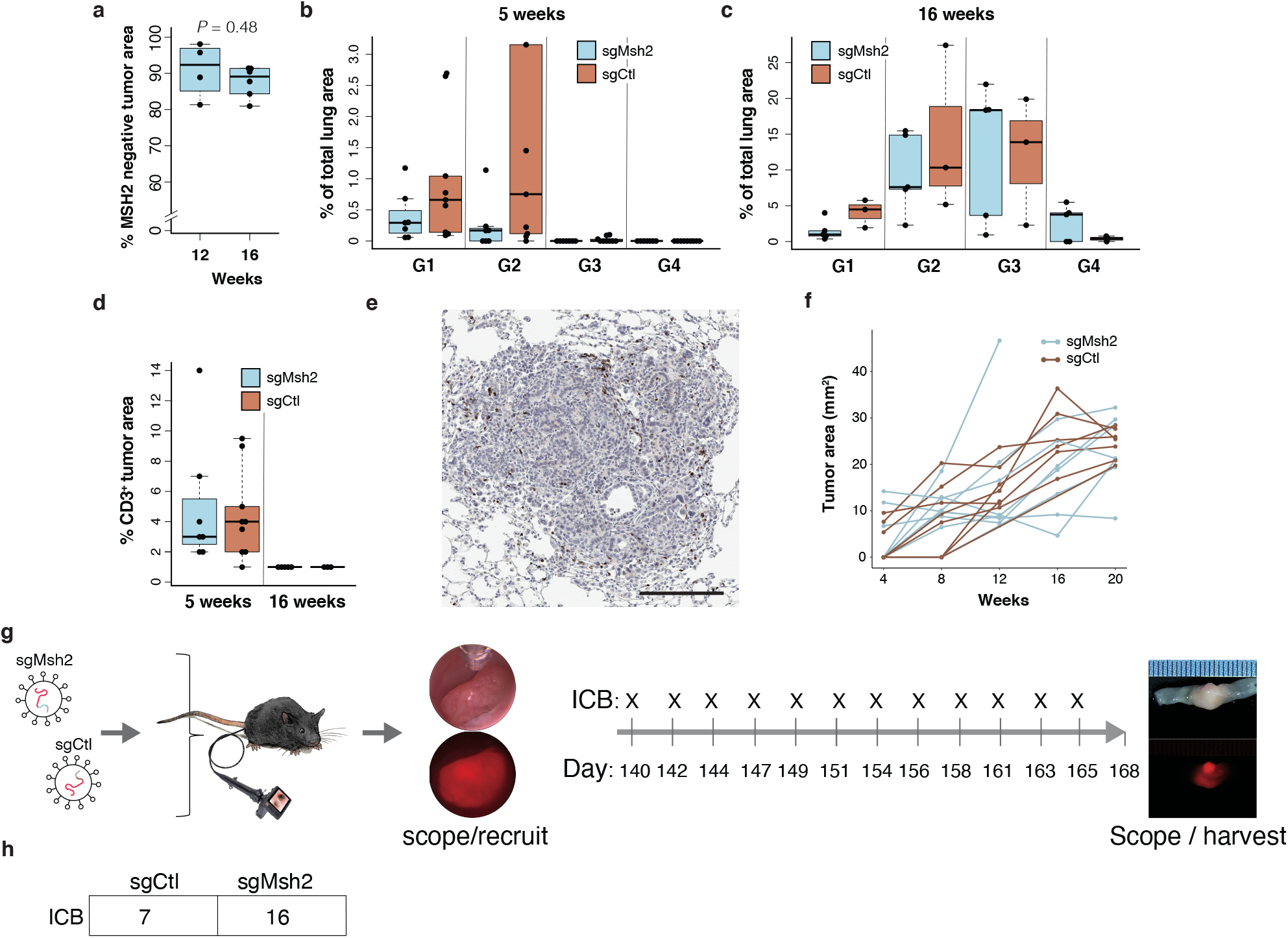
Tumor kinetics and immunogenicity are unaffected by MMRd. **(a)** Percent of overall lung tumor area negative for MSH2 by IHC at 12- and 16-weeks post-initiation of *KP; Msh2^flox/flox^* animals with SPC-Cre adenovirus. N = 4 animals at 12-weeks and 6 animals at 16-weeks. **(b-c)** Percent of total lung area occupied by tumors of grades 1-4 (G1-4) in *KP; R26^LSL-Cas9^* mice at 5- **(b)** and 16-weeks **(c)** post-initiation with sgMsh2- and sgCtl-targeted lentivirus, representative of 7 and 9 animals at 5-weeks and 5 and 3 animals at 16-weeks. Normal lung and tumors were manually annotated and scored in this cohort. **(d)** Average lung tumor area positive for CD3, as a marker of all T cells, by IHC in *KP; R26^LSL-Cas9^* mice at 5- and 16-weeks post-initiation with sgMsh2- and sgCtl-targeted lentivirus. Each point is an independent animal. Number of animals same as in (b-c). **(e)** IHC staining of CD3 in sgMsh2-targeted lung tumor, representative of animals in (d). Scale bar = 200 μM. **(f)** Change in focal colon tumor area over time, as estimated by longitudinal colonoscopy. N = 8 sgMsh2- and 8 sgCtl-targeted animals. **(g-h)** Schematic of colon model preclinical trial design in *R26^LSL-Cas9^* mice targeted with sgMsh2 and sgCtl **(g)**, and number of animals in each group **(h)**. All animals received ICB treatment. Significance in (a) was assessed by Wilcoxon Rank Sum test.

**Supplemental Figure 3.**
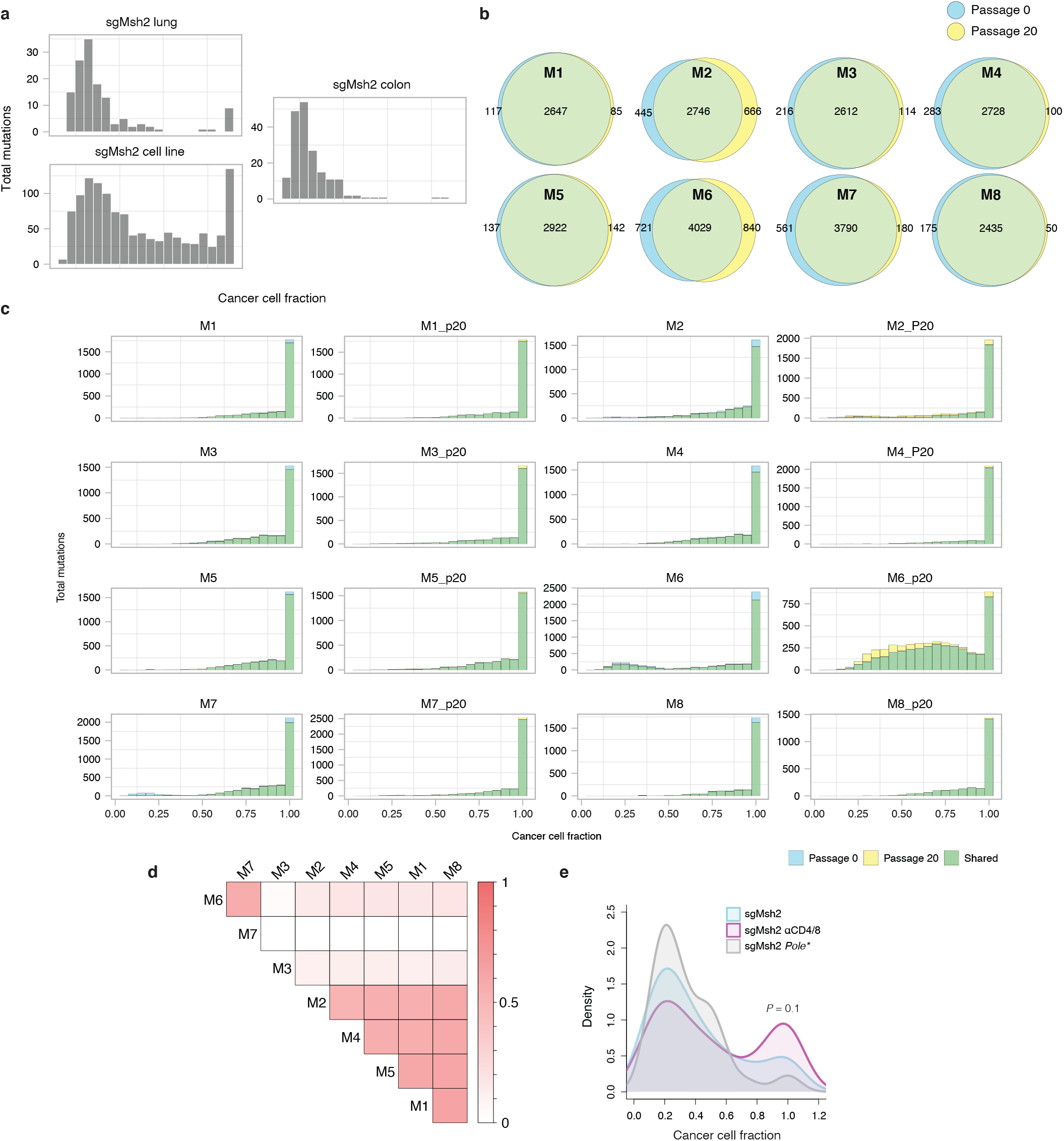
SSCs re-expressing *Msh2* are mutationally stable. **(a)** Histograms of total mutations by cancer cell fraction in a representative sgMsh2-targeted lung tumor, cell line, and colon tumor from Fig. 3a-b. **(b)** Venn diagrams of mutation overlap between M1-8 SSCs sequenced at early passage (called passage 0 for convenience) and 20 passages later. **(c)** Histograms of total mutations by cancer cell fraction in M1-8 SSCs at passage 0 and 20. **(d)** Pairwise intersection map of mutations across M1-8 SSCs. Scale represents fraction of total mutations shared between each pair. **(e)** Distribution of cancer cell fraction estimates of all SNVs in lung tumors from Fig. 3h-j with sgMsh2-targeted *Pole* S415R mutant lung tumor plotted separately (grey). Smoothing was performed by Gaussian kernel density estimation. P-value represents two-sided Kolmogorov-Smirnov test between sgMsh2 and sgMsh2 αCD4/8 distributions.

**Supplemental Figure 4.**
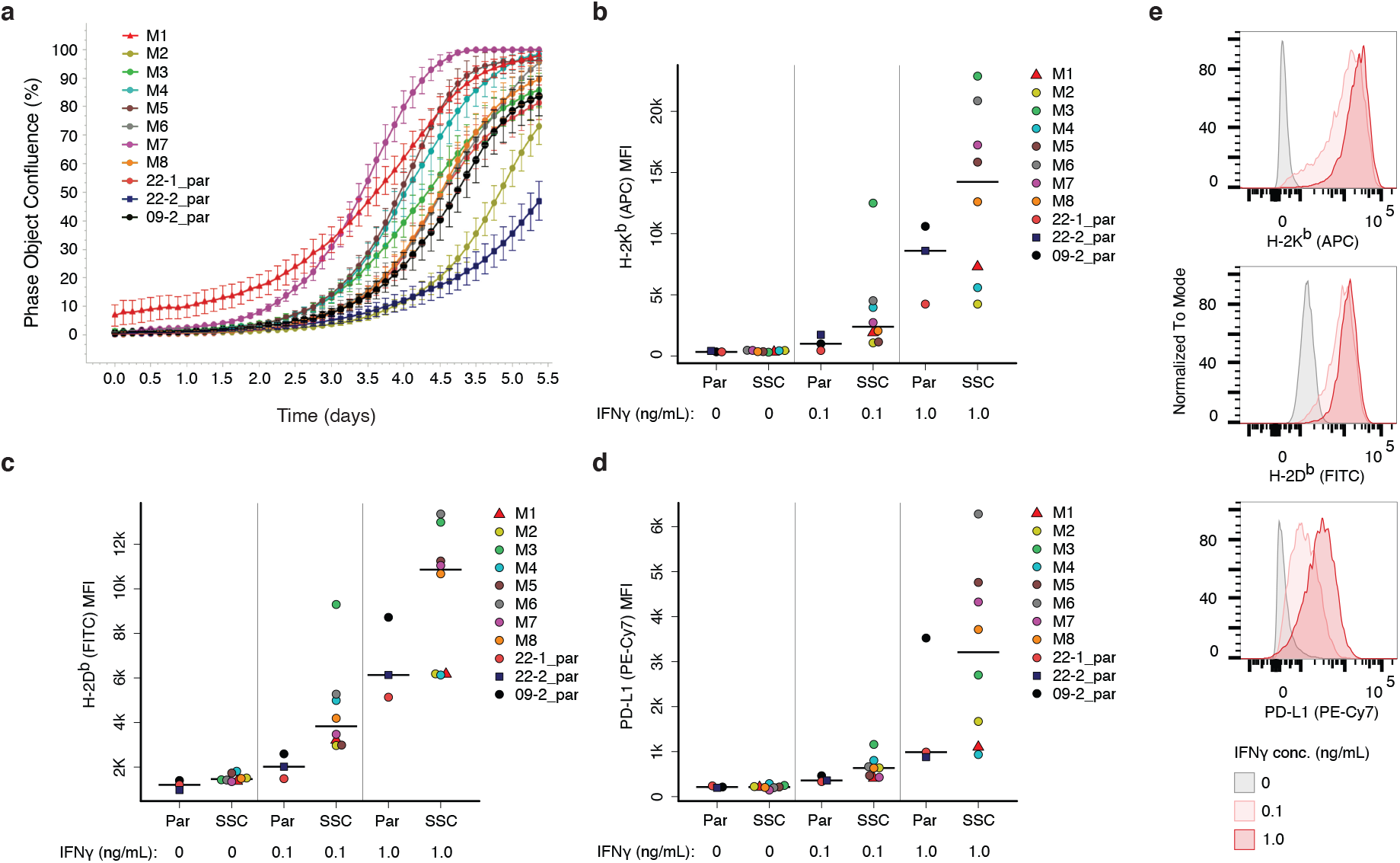
SSCs re-expressing *Msh2* grow similarly *in vitro* and are IFNγ responsive. **(a)** *In vitro* growth kinetics of parental MSH2 knockout cell lines (Par; 09-2, 22-1, 22-2) and *Msh2* re-expressing SSCs generated from 09-2 (M1-8), measured by live cell imaging with an IncuCyte S3. Bars represent standard deviation of eight independent replicates (wells). **(b-e)** Flow cytometric analysis of surface expression of MHC-I alleles H-2K^b^ **(b)** and H-2D^b^ **(c)** and IFNγ-response gene PD-L1 **(d)** in cell lines from (a) following overnight stimulation with 0, 0.1, and 1.0 ng/mL IFNγ. MFI = mean fluorescence intensity. **(e)** Representative histograms of H-2K^b^, H-2D^b^ and PD-L1 expression in an SSC (M3) from the experiment in (b-d).

## Acknowledgements

This work was supported by the NCI Cancer Center Support Grant P30-CA14051, R01 CA233983 and the Howard Hughes Medical Institute. P.M.K.W was supported by a Damon Runyon Fellowship Award. We thank K. Yee, J. Teixeira, K. Anderson, and M. Magendantz for administrative support, and our colleagues in the Jacks laboratory and the broader community at the Koch Institute of MIT for thoughtful discussions and technical advice. We thank the Koch Institute Swanson Biotechnology Center for core support, with particular thanks to K. Cormier and C. Condon for histology support. F.M. and I.C.-C. thank EMBL for funding. We are also grateful for collaboration with T. Westerling and Aiforia in developing automated CNNs for lung tumor grade and IHC quantification.

## Author contributions

P.M.K.W and T.J conceived and directed the study. P.M.K.W, O.S., H.H., N.J.S., A.M.J., W.M.R, and D.A.C. carried out all aspects of the research, animal care and experimentation. F.M. and I.C.-C. designed sequencing analysis pipelines and carried out analysis with P.M.K.W. All other data analysis was carried out by P.M.K.W. Z.A.E. designed and executed the neoantigen prediction pipeline. A.J. provided conceptual and technical guidance in sequencing analysis. S.N. designed dual sgRNA lentiviral constructs. D.Z., M.C.B., A.J., J.J.P., and A.M.C produced and validated critical reagents. P.M.K.W, I.C.-C., and T.J. wrote the manuscript, with feedback from all authors.

## Methods

### Mice

Mice were housed in the animal facility at the Koch Institute for Integrative Cancer Research at MIT with a 12-hour light/12-hour dark cycle with temperatures within 68-72°F and 30-70% humidity. All animal use was approved by the Department of Comparative Medicine (DCM) at MIT and the Institutional Animal Care and Use Committee (IACUC). *Kras^LSL-G12^*^D 51^; *Trp53^flox/flox^* ^52^; *R26^LSL-Cas9^* ^53^ (*KP; R26^LSL-Cas9^*) mice were maintained on an F1 (C57BL/6 x 129/SvJ) background. *Kras^LSL-G12D^*; *Trp53^flox/flox^*; *Msh2^flox/flox^* ^36^ (JAX stock #016231) and *R26^Cas9^* ^54^ (JAX stock # 028555) mice were maintained on a pure C57BL/6 background. Lung cell lines were isolated from tumors induced in albino C57BL/6 hosts chimeric for tissue derived from blastocyst injection of a *KP; R26^LSL-Cas9^* embryonic stem (ES) cell line (12A2) of mixed C57BL/6 and 129/SvJ background and male sex, as previously described^53^. These mice were used in early experiments prior to establishment of a germline *KP; R26^LSL-Cas9^* colony. In orthotopic lung studies, cell lines were transplanted into male chimeras generated from the same 12A2 ES cell line at 10-16 weeks of age. These chimeras are tolerized to C57BL/6 and129/SvJ tissues, potential antigens in the *R26^LSL-Cas9^* allele, and PuroR introduced into cell lines with *Msh2* re-expression (*Kras^LSL-G12D^* tissue in the unrecombined state contains PuroR in the lox-STOP-lox cassette). Autochthonous tumors in lung and colon were induced in approximately equal numbers of male and female mice at 8-16 weeks of age.

### Tumor models

Autochthonous lung tumors in *Kras^LSL-G12D^*; *Trp53^flox/flox^*; *R26^LSL-Cas9^* and *Kras^LSL-G12D^*; *Trp53^flox/flox^*; *Msh2^flox/flox^* mice were induced by intratracheal instillation of 2e4 transduction units (TUs) of lentivirus and 2e8 plaque forming units of adenovirus, respectively, as previously described^30^. Autochthonous colon tumors in *R26^Cas9^* mice were induced by endoscope-guided submucosal injection in the distal colon, as previously described^31,55^. Two injections at 1.5e6 TUs of lentivirus in 50 μL OPTI-MEM were delivered per mouse. Lentivirus was produced in HEK-293 cells (ATCC) and concentrated as previously described^30^, and functional titers (Cre activity, mScarlet fluorescence) measured as previously described^56^. Cell lines were orthotopically transplanted by intratracheal instillation of 1e5 cells in 50 μL SMEM with 5 mM EDTA, followed by a rinse of 30 μL with the same media. Cell lines were established from autochthonous lung tumors by microdissection and mechanical mincing in digestion buffer (HBSS with 1 M HEPES, 125 Units/mL Collagenase Type IV (Worthington) and 20 μg/mL DNAse (Sigma-Aldrich) followed by incubation at 37 °C with gentle agitation for 30 min, and plating in RPMI 10% FBS. Lines were plated into 50:50 RPMI/DMEM 10% FBS at first passage, and DMEM 10% FBS at second passage and thereafter. Cells were taken for WES at third passage. *Msh2* expressing lentivirus was produced as above but without concentration. Cells were incubated with lentiviral supernatant and 3 days later selection begun with 6 μg/mL puromycin (Thermo Fisher), which was maintained during all subsequent culturing.

### Whole-exome sequencing analysis

Whole-exome libraries for all samples were generated using the SureSelect XT Mouse All Exon (Agilent) target enrichment kit with 100 bp paired-end sequencing on the Illumina HiSeq 4000 platform, except for the M1-8 passage 20 SSCs, which were 150 bp paired-end sequencing on the Illumina NovaSeq 6000 S4 platform. Library preparation and sequencing to 100X on-target coverage was performed by Psomagen. Raw sequencing reads were mapped to the GRCm38 build of the mouse reference genome using BWA-MEM v0.7.17-r1188^57^. Aligned reads in BAM format were processed following the Genome Analysis Toolkit (GATK) v4.1.8.0^58^ Best Practices workflow to remove duplicates and recalibrate base quality scores.

SNVs and indels were detected using Mutect2, MuSE v1.0rc^59^, VarDict v1.8.2^60^, and Strelka2 v2.9.2^61^ using matched normal tails as controls. Mutect2 was run using a panel of normals generated using the sequencing data for the 11 tails analysed in this study. Each caller was run independently on each tumor-normal pair, and the calls were integrated using *SomaticCombiner*^62^. For the colon tumors, a panel of four normal tails was used to generate the normal control for all samples, as these mice were of a pure background. We only considered those SNVs detected by Mutect2 and supported by at least one of the other callers. To increase the accuracy of indel detection, only indels detected by at least 2 algorithms were considered for further analysis. Variants mapping to DbSNP (build id 150) positions were discarded. Microsatellite contexts of mutations were annotated using SciRoKo v3.4^63^, with minimum score = 8, seed length = 8, repeats = 2, and mismatch penalty = 1. Only mutations within exons were considered in all analyses.

Somatic copy number aberrations were detected by integrating the output of GATK and FreeBayes v1.3^64^ using PureCN v1.16.0^65^. Briefly, the GATK4 Somatic CNV workflow was utilized for the normalization of read counts and genome segmentation using the tails of all samples to build the panel of normals. FreeBayes was used to obtain B-allele frequency values for dbSNP (build id 150) variant sites of the mouse genome. Finally, PureCN was used to integrate the output of GATK and FreeBayes to estimate the allele-specific consensus copy-number profile, purity, and ploidy of each sample. Ploidy values of cell lines were determined experimentally by metaphase spreads and input into PureCN. Finally, the cancer cell fraction (CCF) value for each SNV was computed using the R package cDriver^40^.

### Clonal deconvolution by targeted amplicon sequencing

To identify private somatic SNVs capable of distinguishing individual SSCs in the parental cell line (09-2_par) and M1-8 SSC mixed tumors (Fig. 3f, 4n), we first selected all clonal SNVs in copy neutral regions of the genome (4 copies, as all lines were tetraploid by metaphase spreads). We then checked the BAM files across all other samples for the presence of reads supporting the alternative allele (base quality > 20 and mapping quality > 30) using an in-house python script relying on the Pysam library. Mutations with support for the alternate allele in at least one sample were discarded. Four private SNVs for each SSC and four common SNVs were chosen for targeted amplicon sequencing, all of which were validated by PCR amplification and Sanger sequencing. 200-250 bp regions spanning these SNVs were individually PCR amplified from samples, gel purified, and submitted for 150 bp paired-end sequencing on the Illumina NovaSeq 6000 S4 platform.

Reads were aligned to a reference fasta file of all targets (+/− 250 nt upstream/downstream of SNV in mouse genome, build GRCm38) using BWA-MEM, with removal of duplicates and recalibration of base quality score following GATK Best Practices. A pileup was then generated using the mpileup function of bcftools v1.10.2^66^ with --min-BQ 30 and a bed file of all SNV coordinates. Using a custom R script, total and SNV-specific depths at all locations were extracted. All SNVs showed more reads supporting the expected alternate allele than other alleles in the M1-8 SSC equal mixture control, except for M6_2, which was excluded from subsequent analysis. Background PCR/sequencing error for each SNV was estimated using the median observed frequencies of SNVs in all metastases (excluding those of the single represented clone in each metastasis), which represented truly clonal controls. SNV frequencies were adjusted by subtracting background values. Clonal percentages in *ex vivo* tumors were estimated by taking the median of private SNV frequencies, multiplying by 4 (SNVs are present on 1 of 4 alleles), and dividing by tumor purity. Tumor purity was estimated by taking the median observed to expected ratio of frequencies of the four common SNVs (present in all SSCs).

### Neoantigen prediction and expression

Variant consequence was annotated using Ensembl Variant Effect Predictor (VEP) v99^67^ with Wildtype and Downstream plugins, the VEP cache and reference genome for GRCm38, and the following parameters: -- symbol, --terms=SO, --cache, --offline, --transcript_version, --pick. The --pick parameter was reordered from default to report the transcript with most extreme consequence for each variant: rank, canonical, appris, tsl, biotype, ccds, length, mane. Neoepitopes were predicted with C57BL/6 mouse MHC-I alleles, *H2-K1* (H-2K^b^) and *H2-D1* (H-2D^b^), and variant effect predictions using pVACtools v1.5.7^46^. Mutant peptides were generated for lengths 8- through 11- amino acids, and MHC:peptide binding affinity was predicted for all peptide:MHC allele pairs with NetMHC-4.0, NetMHCpan-4.0, SMM v1.0, and SMMPMBEC v1.0^42–45^. The median value across all affinity predictions was taken as the final measure of binding affinity.

To assess allele-specific expression of neoantigens, RNA-seq was performed on autochthonous lung tumors (10 sgMsh2 and 10 sgMsh2 with αCD4/8 treatment) and M1-8 SSCs. cDNA libraries were prepared using Kapa mRNA Hyperprep, and 150 bp paired-end sequencing was performed on the Illumina NextSeq platform. Sequencing reads were aligned to the reference genome (GRCm38) using STAR v2.7.1a^68^ and PCR duplicates were removed using Picard v2.23.4^69^. STAR parameters used: outFilterMultimapNmax = 20, alignSJoverhangMin = 8, alignSJDBoverhangMin = 1, outFilterMismatchNmax = 999, outFilterMismatchNoverLmax = 0.1, alignIntronMin = 20, alignIntronMax = 1000000, alignMatesGapMax = 1000000, outFilterScoreMinOverLread = 0.33, outFilterMatchNminOverLread = 0.33, and limitSjdbInsertNsj = 1200000. Considering the somatic variants found in each sample by WES, we used a custom python script to interrogate the presence of these variants in the RNA-seq BAM files. Only non-duplicate reads with mapping quality >= 255 and bases with base-quality >= 20 were considered to compute the variant allele frequencies in RNA.

### Immunohistochemistry

Tissues were fixed in zinc formalin, washed in 70% ethanol and paraffin embedded. Antigen retrieval was performed in citrate buffer pH 6 in a pressure cooker at 125 °C for five minutes. Blocking was performed with BLOXALL Endogenous Peroxidase and Alkaline Phosphatase Blocking Solution (Vector) followed by Normal Horse Serum (2.5%) (Vector). For MSH2 (ab70270, Abcam) and CD3 (ab5690, Abcam) stains, slides were stained overnight at 1:1000, then incubated with HRP anti-Rabbit IgG (Vector) and developed with DAB (Vector). For triple staining (CD8a, CD4, FOXP3), slides were first stained with CD8α (ab217344, Abcam) 1:1000 overnight, incubated with Alkaline Phosphatase (AP) anti-Rabbit IgG (Vector) and developed with Vector Black substrate (Vector). Sections then underwent a second round of antigen retrieval in a pressure cooker at 110 °C for two minutes, followed by co-incubation with FOXP3 (FJK-16s, eBioscience) 1:125 and CD4 (ab183685, Abcam) 1:400 overnight. Sections were then sequentially incubated with AP anti-Rat IgG (Vector) and HRP anti-Rabbit IgG and developed sequentially with Vector Red (Vector) and Vina Green (Biocare Medical). Slides were counterstained with Harris Acidified Hematoxylin and dehydrated. Aqueous wash steps following counterstain were shortened from 1 minute to 30 seconds to minimize loss of Vina Green stain.

### Quantification of tumor burden and immunohistochemistry staining

Quantification of lung tumor burden by grade was performed on scans of hematoxylin and eosin (H&E)-stained sections by an automated deep neural network developed in collaboration with Aiforia Technologies Oy, and in consultation with veterinarian pathologist Dr. Roderick Bronson. A convolutional neural network (CNN) for semantic multi-class segmentation was trained to classify and detect lung parenchyma and non-small cell lung cancer (NSCLC) grades 1-4. For supervised training, selected areas from 93 slides were chosen. The algorithm performed consistently and with high correlation with human graders across multiple validation datasets independent of the training dataset. Version NSCLC_v25 of the algorithm was used.

Immune infiltration in triple stained slides was calculated by a CNN trained to identify the three cell types stained (black = CD8, green = CD4, green/red = Treg), in collaboration with Aiforia Technologies Oy. Whole slides were scanned with a Leica AT2 (Aperio) using the Rainbow color profile. First, the CNN was trained to identify a tissue layer. Within that layer, the CNN was trained to identify black, green, and green/red staining. Within each of these layers, an object counter was trained to quantify the number of cells with the stain. Training was performed by manual annotation of each layer and counting of objects within training regions across 20 separate slides, with roughly five training regions per layer per slide. Performance was validated against human counting and found to be highly accurate and consistent. CD3 infiltration in single stain slide scans was measured as the percentage of pixels positive for stain (DAB) in Aperio ImageScope. Area of positive and negative MSH2 staining was quantified by manual annotation in QuPath v0.1.2^70^.

### Western blot

Cells were lysed in RIPA buffer (Thermo Fisher) supplemented with Halt protease inhibitors (Thermo Fisher) and incubated for 30 min at 4 °C rotating. Protein concentration was determined using Pierce BCA Protein Assay (Thermo Fisher), and equal amounts of protein (20-40 μg) were run on NuPage 4-12% Bis-Tris gradient gels (Thermo Fisher) by SDS-PAGE and transferred to PVDF membranes. Immunoblotting was performed against MSH2 (D24B5, Cell Signaling Technology) at 1:1000, MLH1 (ab92312, Abcam) at 1:1000, GAPDH (6C5, Santa Cruz) at 1:5000, and β-ACTIN (13E5, Cell Signaling Technology) at 1:5000. Blots were stained with HRP anti-Rabbit IgG and developed with Western Lightning Plus-ECL (Perkin Elmer) on X-ray film.

### In vivo antibody and chemotherapy dosing

All antibody dosing was performed via intraperitoneal injection in 100 μl PBS. αCD4 (GK1.5, BioXCell) and αCD8 (2.43, BioXCell) depleting antibodies were administered at 200 ug every 4 days. αPD-1 (29F.1A12, BioXCell) was administered at 200 μg three times a week. αCTLA (9H10, BioXCell) was administered at an initial dose of 200 μg, with all subsequent doses at 100 μg, three times a week. Oxaliplatin (Sigma) and cyclophosphamide (Sigma) (Oxa/Cyc) were co-delivered via intraperitoneal injection in 100 μl PBS at 2.5 mg/kg and 50 mg/kg body weight, respectively, once a week for three weeks, as previously described^39^.

### In vivo tumor imaging and quantification

Lung tumor progression was monitored longitudinally by X-ray microcomputed tomography (μCT) using a GE eXplore CT 120 system, as previously described^71^. Solid lung volume corresponding to tumor burden was quantified using a custom MATLAB (MathWorks) script, as previously described^71^. Colon tumor progression was monitored longitudinally using a Karl Storz colonoscopy system with white light and RFP fluorescence. This consists of Image 1 H3-Z Spies HD Camera System (part TH100), Image 1 HUB CCU (parts TC200, TC300), 175-Watt D-Light Cold Light Source (part 20133701–1), AIDA HD capture system, and fluorescent filters in the RFP and GFP channels (Karl Storz). The endoscope used for imaging was the Hopkins Telescope (Karl Storz, part 64301AA) with operating sheath (Karl Storz, part 64301AA). To consistently measure tumor area, biopsy forceps (Richard Wolf) were fed through the operating sheath and positioned consistently given two landmarks: widthwise grooves that appear as concentric semi-circles in the field of view, and a lengthwise groove at the forceps tip. Images were captured upon gentle contact of forceps with tumor. Tumor area in the field of view and length of the lengthwise forceps groove were calculated using ImageJ v2.1.0/1.53c. Tumor area was normalized to groove length.

### Generation of lentiviral constructs

The previously published U6::sgRNA-EFS::Cre (pUSEC) lentiviral construct^71^ was digested with BsmBI and sgRNAs cloned as previously described^72^. H1::sgApc-U6::sgRNA-EFS::mScarlet was generated by Gibson assembly using an H1::sgApc-scaffold gBlock synthetic gene fragment (IDT), PCR amplicons of U6::BsmBI-filler-BsmBI-scaffold, EFS promoter, and *mScarlet*^73^, and the lentiviral backbone from the Trono laboratory (Addgene). This was then digested with BsmBI and a second sgRNA cloned as above. sgRNA sequences: sgApc: 5’-GTCTGCCATCCCTTCACGTT-3’^31^; sgMsh2: 5’GAACATACATTCGTCAGACCG’; sgMlh1: 5’- GAGGGCACCCTGATCACGGTG-3’; sgMsh3: 5’- CTTACTCCGAGCACTCATCG-3’; sgMsh6: 5’- CATCAGTGACCGTCTAGATG-3’; non-targeting sgCtl (*tdTomato/mScarlet* targeting): 5’- GGCCACGAGTTCGAGATCGA-3’^56^; and targeting sgCtl (*Olfr102*): 5’-GCATCTTTGGCAGTGTCACAG-3’, which were used interchangeably with no observed differences in tumorigenesis. PGK::Msh2-EFS::PuroR was generated by Gibson assembly using multiple gBlocks spanning murine *Msh2* (C57BL/6), PCR amplicons of PGK and EFS promoters and *PuroR*, and the Trono lentiviral backbone.

### Validation of CRISPR/Cas9 editing and estimation of tumor purity

To validate efficiency of gene editing by CRISPR/Cas9, 200-250 bp regions spanning sgRNA sites in the genome were amplified and submitted for deep sequencing (CRISPR sequencing) at the Massachusetts General Hospital DNA Core. Tumor purity in colon tumors was estimated by taking the fraction of edited to wild-type alleles at the sgRNA-targeted site in *Apc* by CRISPR sequencing. Loss of *Apc* is prerequisite for tumorigenesis in the model, and thus an assumption was made that all tumor cells harbor loss-of-function edits at this locus. Tumor purity in sgMsh2-targeted lung tumors was estimated using WES BAM coverage data spanning exons of the *Trp53^flox^* allele and flanking genes (*Wrap53*, *Atp1b2*), which were retrieved using the bedcov function of SAMtools v1.10. The ratio of median coverage in flanking exons (*Wrap53* exons 0-9, *Trp53* exon 11, *Atp1b2* exons 0-6) versus *Trp53* exons flanked by Cre loxP sites (exons 2-10) was calculated in tumors and normal tails. This ratio in tumors, representing extent of *Trp53^flox^* recombination, was then normalized to the median ratio across matched normal tails to estimate tumor purity, with the assumption that all tumor cells, but not normal contamination, underwent complete recombination of *Trp53^flox^* alleles.

### In vitro cell line assays

Serial live cell imaging of cell lines grown in 96-well plates (Corning) and quantification of confluence was performed with an IncuCyte S3 (Sartorius). Eight replicate wells were seeded with 100 cells each per line and imaged every 3 hours for ∼5.5 days. Murine IFNγ (PeproTech) was used for *in vitro* stimulation of cell lines for 24 hours, followed by live/dead staining (ghost ef780 (Corning), 1:500) in PBS and surface staining in 1 mM EDTA, 25 mM HEPES, 0.5% heat-inactivated FBS in PBS with anti-H-2K^b^ (APC, AF6-88.5.5.3, Thermo Fisher), anti-H-2D^b^ (FITC, 28-14-8, Thermo Fisher), and anti-PD-L1 (10F.9G2, PE-Cy7, BioLegend). Samples were run on a BD LSRFortessa using BD FACSDiva v8.0 software. Results were analyzed in FlowJo v10.4.2, excluding dead (ef780 positive) cells.

### Phylogenetic tree

All somatic SNVs and indels called by the WES analysis pipeline in M1-8 SSCs and the 09-2 parental cell line were considered in constructing a phylogenetic tree. Using the R Bioconductor package ‘phangorn’ v2.7.0 and a binary presence/absence matrix of mutations across SSCs and 09-2 as input, a tree rooted on 09-2 was constructed. Specifically, the function prachet was used to calculate the tree using the parsimonious ratchet method, and the function acctran was used to calculate branch length.

### Statistics and reproducibility

Statistical analyses and figure generation were performed in R v4.0.2 using built in functions and ggplot2 v3.3.3, beeswarm v0.3.1, corrplot v0.88, eulerr v6.1.0, and RColorBrewer v1.1.2. For statistical assessment of differences in proportionality, Fisher’s exact 2×2 test was performed. For continuous data, two-tailed Wilcoxon Rank Sum test was performed. Multiple comparison corrections were performed using Holm’s method. No statistical method was used to predetermine sample size. In preclinical trials of lung and colon models, only those animals with apparent tumors by μCT or colonoscopy were recruited. No other data were excluded from analyses. Preclinical trials were randomized, and investigators blinded to allocation during dosing, μCT and colonoscopy imaging, and tumor quantification. No experiments presented in this manuscript failed to replicate.

